# Abnormal local cortical functional connectivity due to interneuron dysmaturation after neonatal intermittent hypoxia

**DOI:** 10.1101/2024.06.04.596449

**Authors:** Ivan Goussakov, Sylvia Synowiec, Rafael Bandeira Fabres, Gabriela Dias Almeida, Silvia Honda Takada, Daniil Aksenov, Alexander Drobyshevsky

**Affiliations:** Department of Pediatrics, NorthShore University HealthSystem, Evanston, IL, USA; Neurogenetics Laboratory, Universidade Federal do ABC, São Bernardo do Campo, Brazil; Department of Radiology, NorthShore University HealthSystem, Evanston, IL, USA

**Keywords:** functional connectivity, resting state fMRI, interneurons, perinatal brain injury

## Abstract

**Background:** Premature infants often experience frequent hypoxic episodes due to immaturity of respiratory control that may result in disturbances of gray and white matter development and long-term cognitive and behavioral abnormalities. We hypothesize that neonatal intermittent hypoxia alters cortical maturation of excitatory and inhibitory circuits that can be detected early with functional MRI.

**Methods:** C57BL/6 mouse pups were exposed to an intermittent hypoxia (IH) regimen consisting of 12 to 20 daily hypoxic episodes of 5% oxygen exposure for 2 min at 37C from P3 to P7, followed by MRI at P12 and electrophysiological recordings in cortical slices and in vivo at several time points between P7 and P13. Behavioral tests were conducted at P41-P50 to assess animal activity and motor learning.

**Results:** Adult mice after neonatal IH exhibited hyperactivity in open field test and impaired motor learning in complex wheel tasks. Patch clamp and evoked field potential electrophysiology revealed increased glutamatergic transmission accompanied by elevation of tonic inhibition.

A decreased synaptic inhibitory drive was evidenced by miniature IPSC frequency on pyramidal cells, multi-unit activity recording in vivo in the motor cortex with selective GABA_A_ receptor inhibitor picrotoxin injection, as well as by the decreased interneuron density at P13. There was also an increased tonic depolarizing effect of picrotoxin after IH on principal cells’ membrane potential on patch clamp and direct current potential in extracellular recordings. The amplitude of low-frequency fluctuation on resting-state fMRI was larger, with a larger increase after picrotoxin injection in the IH group.

**Conclusions:** Increased excitatory glutamatergic transmission, decreased numbers, and activity of inhibitory interneurons after neonatal IH may affect the maturation of connectivity in cortical networks, resulting in long-term cognitive and behavioral changes, including impaired motor learning and hyperactivity. Functional MRI reveals increased intrinsic connectivity in the sensorimotor cortex, suggesting neuronal dysfunction in cortical maturation after neonatal IH. The increased tonic inhibition, presumably due to tonic extrasynaptic GABA receptor drive, may be compensatory to the elevated excitatory glutamatergic transmission.

## INTRODUCTION

Hypoxic perinatal brain injury, a persistent condition despite improvements in neonatal intensive care, can lead to a range of neurodevelopmental impairments, including motor dysfunction, behavioral problems, and cognitive deficits (Dilenge, Majnemer et al. 2001, van Handel, Swaab et al. 2007, Pascal, Govaert et al. 2018). Intermittent hypoxia (IH), a common condition in preterm infants due to immature breathing control (Eichenwald 2016), contributes to impaired neurodevelopment (Pillekamp, Hermann et al. 2007, Poets 2019). Preterm infants, especially those born very premature (<32 weeks gestation), are particularly susceptible to hypoxic brain injury and often exhibit deficits in memory, attention, and executive function (Ketzer, Gallois et al. 2012, Burnett, Scratch et al. 2013, Anderson 2014, Johnson and Marlow 2017, Mikkelsen, Olsen et al. 2017).

Animal models of perinatal hypoxia reproduce cognitive and behavioral deficits reminiscent of human conditions, including learning, memory, executive function impairments, hyperactivity, and attention problems (Sab, Ferraz et al. 2013, Maxwell, Zimmerman et al. 2020). These findings suggest that perinatal hypoxia disrupts the delicate balance between excitatory and inhibitory circuits within the developing brain. Studies report a decrease in major interneuron types (Komitova, Xenos et al. 2013, Chavez-Valdez, Emerson et al. 2018, Stolp, Fleiss et al. 2019, Yang, Davidson et al. 2023). However, the functional impact of this interneuron injury on neuronal network activity and glutamatergic transmission remains unclear. Mice exposed to neonatal hypoxia exhibit increased neuronal activity and are prone to seizures (Burnsed, Skwarzyńska et al. 2019). This elevated excitability is attributed to increased AMPA receptor transmission (Jensen, Wang et al. 1998, Lippman-Bell, Zhou et al. 2016). In a mouse neonatal IH model, we also observed abnormalities in glutamatergic transmission, including elevated NMDA and AMPA currents, along with a reduction in AMPA-lacking silent synapses (Goussakov, Synowiec et al. 2021). Disruptions in the excitatory/inhibitory balance, caused by increased excitatory transmission and decreased interneuron function after perinatal hypoxia, can impair synaptic pruning and neuronal circuit formation during this critical developmental period (Rakic, Bourgeois et al. 1986). This can lead to hyper-connected networks (Kerchner and Nicoll 2008), potentially contributing to cognitive, behavioral, and memory deficits. Additionally, the maturation of principal cells and interneurons is influenced by the shift of GABAergic transmission from excitatory to inhibitory (Sauer and Bartos 2010, Ben-Ari, Khalilov et al. 2012). A crucial question remains whether functionally deficient interneurons provide sufficient inhibitory modulation to counteract the increased glutamatergic transmission.

We hypothesized that functional injury to interneurons after neonatal global hypoxia disrupts cortical maturation of the excitatory/inhibitory balance at the level of synaptic and extra-synaptic phasic and tonic modulation of glutamatergic transmission, leading to altered long-term cognitive and behavioral outcomes. We focused on the motor cortex because sensory-motor abnormalities are a major clinical manifestation of perinatal brain injury, including impairments in motor learning and voluntary control (Cahill-Rowley and Rose 2014), as well as motor hyperexcitability and hyperactivity (van Handel, Swaab et al. 2007). This study investigated the feasibility of pharmacological functional MRI as a non-invasive and potentially clinically valuable tool for the early detection of interneuron-specific abnormalities in cortical networks following neonatal hypoxia.

## METHODS

The study has received approval from the Institutional Animal Care and Use Committees of NorthShore University HealthSystem.

### Neonatal intermittent hypoxia

C57BL/6J mice were originally obtained from the Jackson Laboratory (Bar Harbor, Maine) and bred at the NorthShore University HealthSystem animal facility. Neonatal mice of both genders were randomly assigned to the intermittent hypoxia (IH) or control group. Animals underwent intermittent hypoxia using a gas mixture of 5% O2 and 95% N2 for 2.5 minutes, followed by 5 minutes of re-oxygenation (Goussakov, Synowiec et al. 2019). During hypoxic exposure, pups were separated from their dams and placed into a 250 mL air-tight chamber in a temperature-controlled neonatal incubator with an ambient temperature of 35 °C. The paradigm began at postnatal day 3 (P3) and continued for 5 consecutive days. Airflow was maintained at 4 L/min and switched between the hypoxic mixture and room air using solenoid valves controlled by a programmable timer. Oxygen concentration and air temperature within the chamber were continuously monitored with calibrated sensors, reaching a nadir of 5% within 40 seconds of initiating the hypoxic period. The total number of daily hypoxic episodes gradually decreased across the 5 days (20, 18, 11, 9, and 8), divided into morning and afternoon sessions. Between sessions, pups were returned to their dams for 4 hours for recovery and nursing. Control mice underwent similar handling procedures, but remained in room air within the incubator throughout the experiment.

### Motor cortex slice preparation and patch clamp recording of Betz cells

Mice between P50-P60 were deeply anesthetized with isoflurane (3%) and decapitated.

Brains were rapidly removed in ice-cold artificial cerebrospinal fluid (aCSF) solution and the 300 µm thick slices, containing motor M1 cortex, were obtained at the level of forelimbs according to Paxinos atlas (Paxinos and Franklin 2004) using Leica vibrotome (VT1000S, Leica Biosystems). The aCSF solution for slice preparation consisted of the following reagents in mM: 68 NMDG, 2.5 KCl, 1.25 NaH_2_PO4, 30 NaHCO_3_, 20 HEPES, 2 Thiourea, 25 D-glucose, 3 Na-ascorbate, 0.3 ml/L Ethyl Pyruvate, 0.5 CaCl_2_-2H2O, 10 MgSO_4_-7H_2_O, 2 Kynurenic acid. The final pH was adjusted to 7.3 with 1N HCl, osmolality was 300±5 mOsm/kg H_2_O. The NMDG was used for slicing as a substitute for NaCl, to increase cell viability in slice preparation. Slices were transferred to a custom-made incubation chamber with a storage solution, containing in mM: 120 NaCl, 2.5 KCl, 1.25 NaH_2_PO_4_, 25NaHCO_3_, 10 D-glucose, 2 Thiourea, 3Na-ascorbate, 0.3 ml/L Na-Pyruvate, 2 CaCl_2_-4H2O, 2 MgSO4-7H_2_O, 2 Kynurenic Acid and10 HEPES/NaOH, pH 7.3, and osmolality 300±5 mOsm. Slices were maintained on nylon mesh in an incubation chamber and bubbled with carbogen for at least 1 hour before recording. For patch-clamp recording slices were transferred to a submerged chamber (model RC-24, Warner Instruments Hamden, CT, US) and perfused at a rate 3 ml/min at room temperature with aCSF of following composition in mM: 120 NaCl, 2.5 KCl, 25 NaHCO_3_, 2 CaCl_2_, 2 MgSO_4_, 1.25 NaH_2_PO4, 10 HEPES, 10 D-glucose, bubbled with carbogen (95% O_2_/5% CO_2_) to maintain pH of 7.4. Electrodes were pulled from capillary tubes 1B150F-4 (World Precision Instruments Inc., FL) using P97 puller (Sutter Instrument, CA) and filled with a solution composed in mM: 110 K-Gluconate, 15 Cs-methanesulfonate, 10 TEA-Cl, 5 QX-314, 4 NaCl, 2 MgCl_2_, 10 EGTA, 10 HEPES, 2 MgATP, 0.3 NaGTP for miniature currents recording in whole cell configuration. The solution was titrated with CsOH to pH 7.2 with an osmolality of 290±5 mOsm. Filled electrodes had a resistance of 2–5 MΩ. Pipette capacitance and whole-cell compensations were applied.

For patch clamp monitoring of transmembrane potential (Vm), Cs-methanesulfonate was not used to avoid a partial block of chloride ion channels.

Betz cells, identified by the size of 30-40 μm and location in L-V of the primary motor cortex, were recorded in whole-cell configuration. Series resistance (Rs) was not compensated but monitored and recordings with more than 15% change in Rs were discarded. Post-synaptic currents (PSCs) were amplified using Axopatch-700B amplifier, digitized at the sampling rate of 100 kHz, and recorded in a voltage clamp using pCLAMP-10 software (Molecular Devices, US). Liquid junction potential was calculated using a tool in PClamp10 as -9.8 mV and resting membrane potential (Vm) was corrected in data analysis. Vm was measured in the current clamp mode. Spontaneous miniature currents events were isolated by blocking action potentials with 1 μm tetrodotoxin (TTX). Glutamatergic miniature currents (EPSCs) were, measured at holding potential of −70 mV as inwardly directed currents. After 10 min recording time, the holding potential was changed to 0 mV, and spontaneous GABAergic postsynaptic currents (IPSCs) were recorded for another 10 min as outwardly directed currents. The polarity switch sequence was altered randomly between recordings. In control experiments, the glutamatergic nature of EPSCs was confirmed by a complete block of spontaneous activity with 6-cyano-7-nitroquinoxaline-2,3-dione disodium salt (CNQX, 50 μM, Sigma-Aldridge, US) and D-2-amino-5-posphonopentanoic acid (APV, 100 μM; Tocris, US) at a holding potential−70 mV (n = 12).

Spontaneous IPSCs were identified as GABAergic by their suppression with 100 μM picrotoxin (PTX, Tocris, USA) at 0 mV holding potential. PSCs were analyzed with MiniAnalysis software (version 6.0.3 (Synaptosoft, Decatur, GA, USA) by manual selection events over a 2.5 pA threshold. At least 200 of each PSC’s per cell was selected for analysis. Mean amplitudes and frequency of PSC were calculated. Tonic GABAA receptor inhibition was explored by monitoring Vm in whole-cell configuration during 20 min perfusion of Picrotoxin, a specific inhibitor of these receptors.

### Field potential recording and analysis

Field potential recording was performed at multiple age points in mouse pups between P7-P13 and in adults at P50-P55. Brains were removed into ice-cold aCSF, which contained a small amount of ice to keep the temperature at around 2°C. The composition of the preparation, incubation, and recording of aCSF was the same as described above for patch clamp recording preparation. The frontal half of the brains were separated and sliced on a vibratome VT1000S (Leica Biosystems, Deer Park, IL) at 300 µm thickness at the area around the M1 location and subsequently placed on nylon mesh in a storage chamber. After incubation for at least one hour, the slices were transferred to the submerged recording chamber model RC-24 (Warner Instruments, Hamden, CT, US) and perfused at a rate of 3 ml/min by gravity. At this rate, the 500 µl volume of the recording chamber is refilled in 10 seconds. A glass recording electrode, filled with aCSF was placed in the middle of the molecular layer of the M1 cortex. A bipolar stimulation electrode, composed of tightly twisted 25 µm diameter platinum/iridium wires, was placed 200-400 µm away from the recording site in the middle of the molecular layer. The distance between the electrodes was manipulated to avoid pop-spikes that may otherwise introduce secondary activation by firing stimulated afferent cells at all ranges of stimulation intensity 0-700 µA, All recordings were made with a patch-clamp amplifier Multiclamp 700B (Molecular Devices) with low-pass filter disabled to measure DCP from the sweep baseline. The high-pass filter of the amplifier was set to 5 kHz, and the Digidata 1242 was used to sample at 50 kHz and run by pClamp 10.7 software (Molecular Devices).

A 100 µs duration monopolar pulse stimulus from a stimulus isolator A360 (World Precision Instruments, Sarasota, FL), triggered from Digidata 1442 by Clampex 10.1 software. Input-output curves of the monosynaptic field excitatory postsynaptic potential (fEPSP) response were recorded in normal aCSF in a range of stimulus intensity 0-700 µA, with a step of 50 µA and once per minute repetition, 2-6 averages. Baseline fEPSP response with constant stimulus intensity to 600 µA pulse was recorded in aCSF for at least 20 minutes until a stable baseline was achieved during at least 15 minutes, followed by at least 15 minutes (or as long as it was needed to get a steady state plateau on the FEPSP amplitude) recording with 200 µM Picrotoxin (PTX) in perfusate. The stimulus intensity of 500 µA was chosen to evoke a reliable FEPSP in the neonatal cortex without a pop spike appearance The 200 µM PTX concentration was selected after several pilot recordings to produce a substantial effect in neonatal mice cortex without invoking seizure activity. The swipes were recorded in Clampex for all time between triggering stimulus isolator to monitor possible seizure activity during PTX perfusion.

The initial slope of fEPSP was measured within 3 ms latency. The time course of the PTX effect was calculated as a percentage of the average baseline, recorded in normal aCSF. Direct current potential (DCP) was measured as the averaged pre-stimulus sweep baseline magnitude to evaluate tonic GABA receptors mediated inhibition with the addition of PTX. A total of 2-4 slices was recorded per animal. The recordings were averaged per slice and then averaged per each animal for final comparison.

### Animal instrumentation for in vivo MRI and electrophysiological recording

At P11 mouse pups were instrumented with a custom 3D printed circular head post 10 mm diameter with an opening 6 mm diameter (Supplementary Figure 1) with a small height profile to minimize disturbances to the dam while nursing in the nest. During MR imaging and electrophysiological recording, the head post was coupled with an adapter allowing head fixation to the animal cradle. The head post was attached to the skull using acrylic adhesive and dental cement. A small cranial window of about 2x2 mm was made using a 25G needle above the right motor cortex with coordinates AP 1.5 ML 1.0. A plastic cannula and electrode guides were positioned above the cortex and the opening was covered with dental cement. Animals received post-op analgesic Buprenex 0.1 mg/kg and left to recover at least 24 hours before imaging and electrophysiological recordings.

### Resting-state pharmacological functional MRI

Resting-state pharmacological functional MRI was performed on a 9.4 T Bruker Biospec system (Bruker, Billerica, MA) in mouse pups at P12. Mice were sedated with isoflurane (Abbot, IL) inhalation, diluted in air to 3% for induction and 1.0% for maintenance. The animal’s respiration rate and rectal temperature were monitored with a small animal physiological monitor (Model 1030, Small Animal Instruments, NY, USA). Body temperature was monitored using a rectal probe and kept at 35C by blowing warm air. Animals were positioned prone in a cradle with their heads secured beneath the surface coil. The cradles were then inserted into the scanner bore and aligned against a stopper located on the cradle rails. The cradle retraction and repositioning system achieved a sub-millimeter accuracy of approximately 0.3 mm along the scanner’s z-axis. The receiver coil was a 16 mm surface coil (Doty Scientific, SC, USA) allowing full mouse brain coverage. The transmitter was a 70 mm quadrature volume coil. The anesthesia was discontinued to allow animals to wake up for at least 30 min before functional imaging. Anatomical reference was acquired using RARE sequence TR/TE/NEX 3800/14/2 covering the cerebral cortex. Resting-state functional MRI (rsfMRI) time series with the same geometry as the reference image was acquired using single-shot gradient echo T2*-weighted EPI, TE 18 ms, temporal resolution 1.5 sec between volumes, 400 volumes. The field of view was 2.0x1.5 cm, matrix 128x96, in-plane resolution 0.156x0.156 mm^2^, slice thickness 0.5 mm. After the first fMRI run, the cradle was retracted and 1 µL of PTX, 0.1 mg/mL was slowly injected into the motor cortex through the cannula guide on the depth 1 mm using a micro-injector and Hamilton syringe with 33G needle during 5 min and left in the tissue for additional 1 min before slow retraction. The injection site was marked with a small bead of petroleum jelly to aid the detection of MR images. The cradle was reinserted inside the scanner against the positional stopper. A new reference scan was acquired. The second rsfMRI run was performed 15 min after the PTX injection.

Post-processing will be done with custom scripts in Matlab. Brain volumes were motion-corrected using SPM12 and 9 degrees-of-freedom affine transformation. Severe motion outliers were detected and removed using the ArtRepair toolbox for SPM (https://www.nitrc.org/projects/art_repair). No spatial filtering was applied. Linear regression analysis will be performed to remove sources of spurious variance using nuisance variables: 6 parameters obtained by rigid body analysis of head motion and a signal from cerebrospinal fluid regions of interest located in the lateral ventricle.

Maps of Amplitude of Low-Frequency Fluctuations (ALFF), fractional ALFF (fALFF), and regional homogeneity (ReHo) were calculated with a Matlab routine modified from the REST toolkit (https://www.nitrc.org/projects/rest/). The signal in each voxel was normalized to a global mean (Xi, Zhao et al. 2012). After transforming voxel time series frequency information into the power domain, ALFF was calculated as the sum of amplitudes within a low-frequency range (0.008-0.3Hz) [20]. fALFF was intended to reduce the sensitivity of ALFF to physiological noise by taking the ratio of the power spectrum in each frequency band (0.008-0.1 Hz, (0.1-0.3Hz) to the total frequency range (0-0.3Hz). ReHo [21] is defined as the connectivity of a given voxel to those of its nearest 19 neighboring voxels using Kendall’s coefficient of concordance. A 1.5x1 mm oval region of interest (ROI) was placed around the injection site and mean values of the above rsfMRI measures were extracted from the fMRI runs before and after the PTX injection. The analysis was conducted in the native imaging space.

### Extracellular multi-unit recording in awake neonatal mice

P13-P14 mice were sedated with 1.5-2% isoflurane and placed in a custom cradle with the head secured using a head post adapter. The animal was lightly swaddled in gauze to maintain body temperature and minimize movement. The cradle was then positioned in a stereotaxic device. Following anesthesia termination, a 20-minute wait period ensured the animal was fully awake before starting the recordings (Aksenov, Dmitriev et al. 2018). A 12 MΩ tungsten electrode (A-M Systems, Inc., USA, cat # 577200) was lowered through the implanted guide cannula targeting layer 4 of the motor cortex at stereotaxic coordinates AP 1.5, ML 1. A ground wire was placed subcutaneously in the neck region. Electrophysiological signals were amplified and filtered through a miniature preamplifier connected to a multi-channel differential amplifier system (Neuralynx, Inc., Bozeman, MT, USA). The signals were amplified, band-pass filtered (300 Hz-3 kHz for multi-unit activity (MUA)), and digitized at 32 kHz per channel using a Neuralynx data acquisition system. Offline MUA analysis was performed using threshold detection followed by waveform analysis to exclude noise, using software such as SpikeSort 3D (Neuralynx, Inc., Bozeman, MT, USA). The threshold criteria were set to 3 times the estimated standard deviation of the background noise (Aksenov, Li et al. 2023). Following a 9-minute baseline recording, 1 µL of PTX solution (0.1 mg/mL in artificial cerebrospinal fluid (ACSF)) was superfused through the cannula guide above the recording area. This specific concentration of PTX was chosen based on previous and preliminary studies demonstrating sufficient changes in neuronal activity without inducing epileptiform activity in the recordings of healthy animals depending on the method of infusion (Aksenov, Li et al. 2023). Control animals received equal volumes of vehicle (ACSF) using the same procedure to account for potential injection-related effects. An additional 30 minutes of electrophysiological recordings were obtained following PTX or ACSF injection. Finally, the animals were euthanized.

To assess the effects of GABAergic inhibition, our analysis focused on comparing multi-unit activity (MUA) within and between groups. First, we compared MUA values before and after PTX/ACSF injection within each group (control and PTX). Second, we compared MUA values between the control and PTX groups. Since the peak effect of PTX might occur at slightly different times due to variations in drug diffusion across animals, we segmented the post-injection data into 5-minute intervals. We then exclusively utilized the 5-minute interval within each animal that exhibited the maximum change in MUA for our statistical comparisons. The analysis of the control group data also incorporated the time window with the maximum change in MUA for consistency. To assess changes within each group induced by the injection, we utilized a paired two-tailed t-test. This non-parametric test is particularly suitable as it analyzes repeated measures from the same animals. For comparisons between groups (control vs. PTX), we used a two-tailed, two-sample unequal variance t-test. This test is suitable when comparing groups with potentially unequal variances in MUA.

### Behavioral assessment

All behavioral testing procedures were conducted in a dedicated quiet room between 41 and 50 days of age during the daytime. Each apparatus was thoroughly cleaned with 70% ethanol and aired for 3 minutes between animals. Animal movements were tracked and analyzed using ANY-maze software version 7.3 (Stoelting Co., Wood Dale, IL, USA).

### Open Field Test

Mice were placed individually in the center of a clear open field box (61 x 61 cm) and allowed to explore freely for 20 minutes. The software automatically recorded the following parameters of spontaneous locomotor activity: total mobile time, mean speed, total distance traveled, and time spent in the center zone (40 x 40 cm) compared to the peripheral zone. The center zone time is often used as an indicator of anxiety-like behavior in rodents.

### Accelerating Rotarod Test

This test assessed balance, motor coordination, and motor learning. Mice were placed on a horizontal rotating rod set at 4 rpm (Harvard Apparatus, Holliston, MA). The rod’s speed gradually accelerated from 4 to 40 rpm over a 5-minute trial. The latency to fall (time spent on the rod) was recorded. Each mouse received 4 trials per day for 5 consecutive days, with 20-60 minutes of rest in their home cage between trials.

### Complex Wheel Test

A training wheel with all 38 rungs (Lafayette Instrument, Lafayette, IN) was presented individually for 3 consecutive days during day time allowing for normalization of running behavior. On the fourth day, the standard wheel was replaced with a complex wheel of the same diameter but with 22 rungs missing in an alternating pattern. Running activity was recorded for an additional 4 days. Wheel speed was measured using an optical sensor and custom software with a 1-second resolution. The session duration was adjusted to ensure 1 hour of active running time for each animal. The following outcome measures were calculated to assess motor performance and learning: total distance traveled, maximum and average speed, duration of running episodes, and average variability in running speed (coefficient of variation) in 10-second bins.

### RNA Isolation and Real-time PCR

Mouse brains were collected, motor cortex was separated and individually frozen on liquid nitrogen and stored at -80°C before use. The tissue was homogenized in Qiazol Lysis Reagent (cat. # 79306 Qiagen, Germantown, MD) on ice for extraction of total RNA. NanoDrop (ThermoFisher Scientific, USA) instrument was used to measure RNA quantity and quality. 500 ng of isolated total RNA was used to synthesize cDNA using an RT2 First Strand Kit from QIAGEN. cDNA was amplified by polymerase chain reaction (PCR) with QuantiTect Sybr Green PCR Kit (Qiagen, Germantown, MD) on an Applied Biosystems QuantStudio 7 Flex real-time quantitative PCR instrument. The total reaction volume of 25 μL consists of 1.0 µl RT product (cDNA). Primers for Gria1-4, Gad1, Gad2, Slc12a2, and Slc12a5 GAPDH for Sybr Green Assay were predesigned and ordered from Integrated DNA Technologies, Inc., USA. The primer sequences are listed in Supplementary Table 1. For each primer, the final concentration was 0.4 μm and the annealing temperature was adjusted. Gene expression was normalized to the housekeeping gene GAPDH and expressed as a fold change of experimental controls ΔΔCt method.

### Stereological assessment of interneurons subpopulations

Mice were euthanized at P12 and P60 by transcardial perfusion with saline solution (0.9% NaCl) followed by 4% paraformaldehyde (PFA). Brains were removed and post-fixed in 4% PFA overnight, soaked in 30% sucrose solution for cryoprotection, frozen on dry ice, and cryosectioned 20 μm in the coronal plane. Primary antibodies against parvalbumin (ab181086, Abcam), and somatostatin (ab111912, Abcam) were used correspondingly. This was followed by incubation with biotinylated secondary goat anti-rabbit IgG antibodies (1:200; Vector, Burlingame, CA, USA) for 1 hour at room temperature and Avidin biotin complex for 1 hour. The color was developed using 3,3′-diaminobenzidine (Sigma Aldrich). Stereological estimation of neuronal cell density was performed under a microscope (Leica Microsystems, Wetzler, Germany) attached to a motorized stage (Ludl Electronic Products, Hawthorne, NY). Optical fractionator probes were used to estimate the number of labeled cells in the motor cortex in Stereo-Investigator software (MBF Bioscience, Williston, VT, USA). Six 20 μm -thick coronal sections 300 μm apart, starting approximately 1 mm from the end of the olfactory bulbs and covering primary and secondary motor cortex, were counted with a counting frame 350 × 350 μm^2^ and a sampling grid of 1100 × 1100 μm^2^. The coefficient of error of the stereological estimation (Gundersen) for each animal ranged from 0.05 to 0.1. Cell density was estimated by dividing the estimated cell number by the measured tissue volume.

### Statistical analysis

For comparisons of normally distributed data between the two groups, we employed a two-tailed t-test. To assess the main and interaction effects of postnatal age, sex, and IH exposure, we performed two-factor ANOVAs. Post-hoc Sidak’s multiple comparisons test was then used to identify specific group differences revealed by the ANOVA. Repeated-measures (RM) ANOVA was employed to analyze multi-day behavioral data and the effects of drug injections within subjects. Statistical analyses were performed using MATLAB R2023b (MathWorks, Natick, MA) and GraphPad Prism 10 (GraphPad Software, Inc., La Jolla, CA). A significance level of p ≤ 0.05 was used.

## RESULTS

### Hyperactivity and impaired motor learning in adult mice after neonatal IH

Adult mice after neonatal IH were hyperactive with increased locomotion speed, mobile time, and traveled distance (Figure 1A). A two-way ANOVA was performed to evaluate the effects of sex and IH on traveled distance. The results indicated a significant main effect for IH, F (1, 33) = 25.33, p = .0025. The main effect of mouse sex and interaction between sex and IH were not significant. No significant effects were found for the time in the center zone, recorded as a measure of anxiety (Figure 1B).

**Figure 1.**
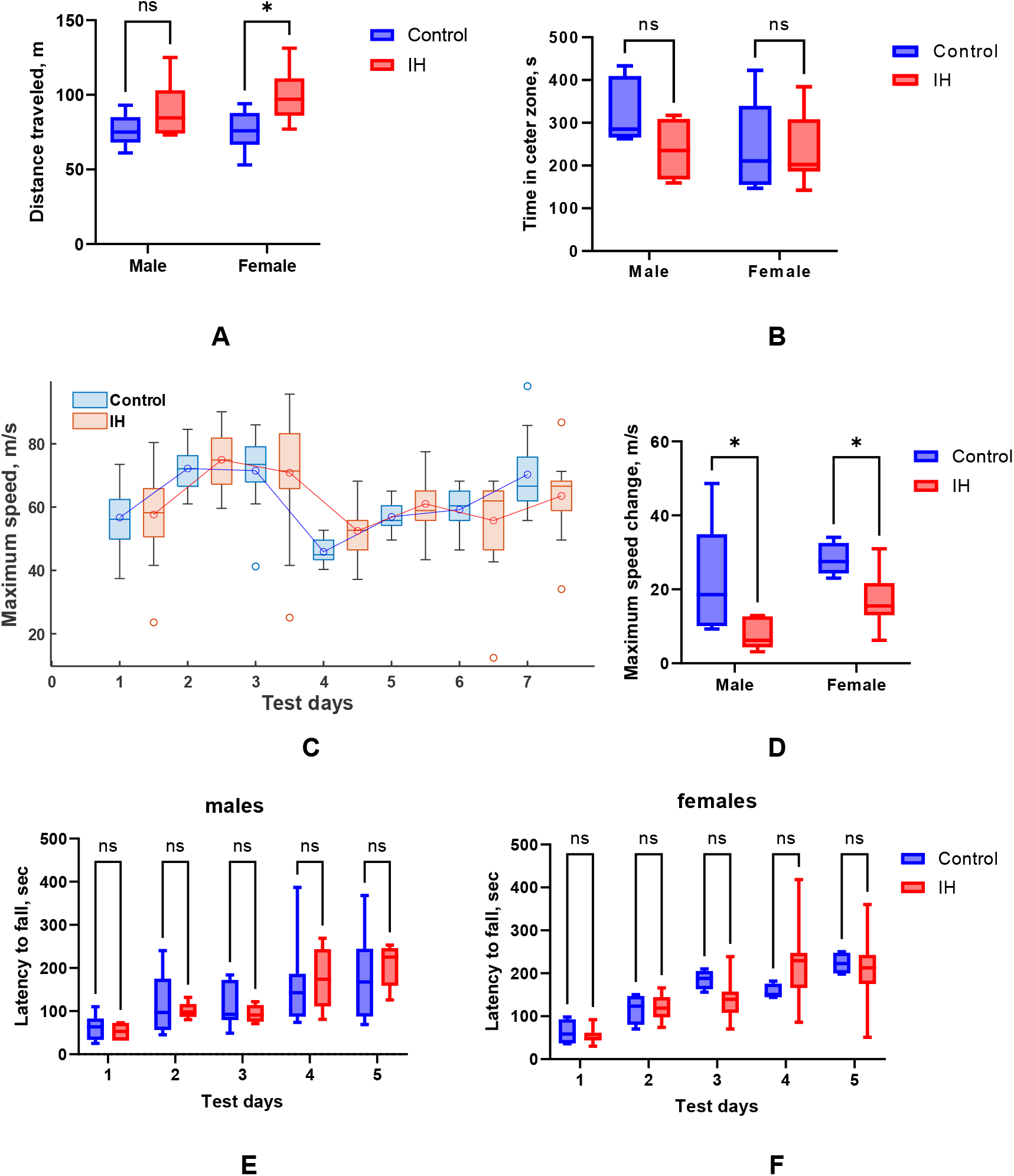
Tests for motor activity and motor learning. Increased locomotion in the open field test (A) was observed in females in IH group, but was not different in time spent in the open field central zone (B). Maximum speed in complex wheel (C) was similar in full wheel days 1-3 between Control and IH groups, but significantly changed in complex wheel days 4-7. The increase in the maximum speed between days 4 to 7 was less in IH group (D). E, F – Rotarod test. Latency to fall increase with test days was but not different between the groups. *-p<0.05, ANOVA post-hoc test.

Motor learning was assessed with a complex wheel test. When a full wheel was presented to individual mice for 3 consecutive days, the maximum running speed increased and reached the plateau by day 3 in both the control and IH groups (Figure 1C). However, in the complex wheel with missing runs at day 4, the maximum running speed decreased to a larger degree in the control group, but was larger than in the IH group by day 7. The interaction term between IH and testing day on maximum running speed was significant on RM ANOVA (F (3, 72) = 4.067, p=0.01). A two-way ANOVA was performed to evaluate the effects of sex and IH on the change in maximum running speed between days 4 and 7, with both sex F (1, 26) = 4.269, p=0.048)and IH (F (1, 26) = 0.3821, p=0.0008) factors been significant. Intriguingly, IH mice displayed faster running speeds in the complex wheel on day 4. However, video recordings revealed a larger number of missteps in this group, suggesting potential motor incoordination despite the increased speed.

Accelerating rotarod, performed for 6 consecutive days, revealed successful motor leaning, indicated in increased time to fall (test time main effect F (1.928, 44.34) = 40.16, p<0.0001) but was not different between control and IH groups either for males and females (Figure 1 E, F).

### Decreased inhibitory drive and increased excitatory drive on Betz cells in the motor cortex of adult mice after neonatal IH

Patch clamp recording in pyramidal Betz cells of the primary motor cortex was performed on a subset of adult mice after neonatal IH at -70 mV holding potential to examine spontaneous excitatory glutamatergic postsynaptic currents (Figure 2 A, B) and at 0 mV holding potential to examine spontaneous GABAergic postsynaptic currents (Figure 2 C, D). The amplitudes of EPSCs were higher in the IH group (t=8.001, df=26, p<0.0001), but the EPSC’s frequency was not different from the control group *Figure 2 E, F). For IPSCs (Figure 2 G, H), the amplitudes were not different between the control group and IH groups, but the IPSC’s frequency was lower in the IH group (t=3.202, df=16, p=0.0056).

**Figure 2.**
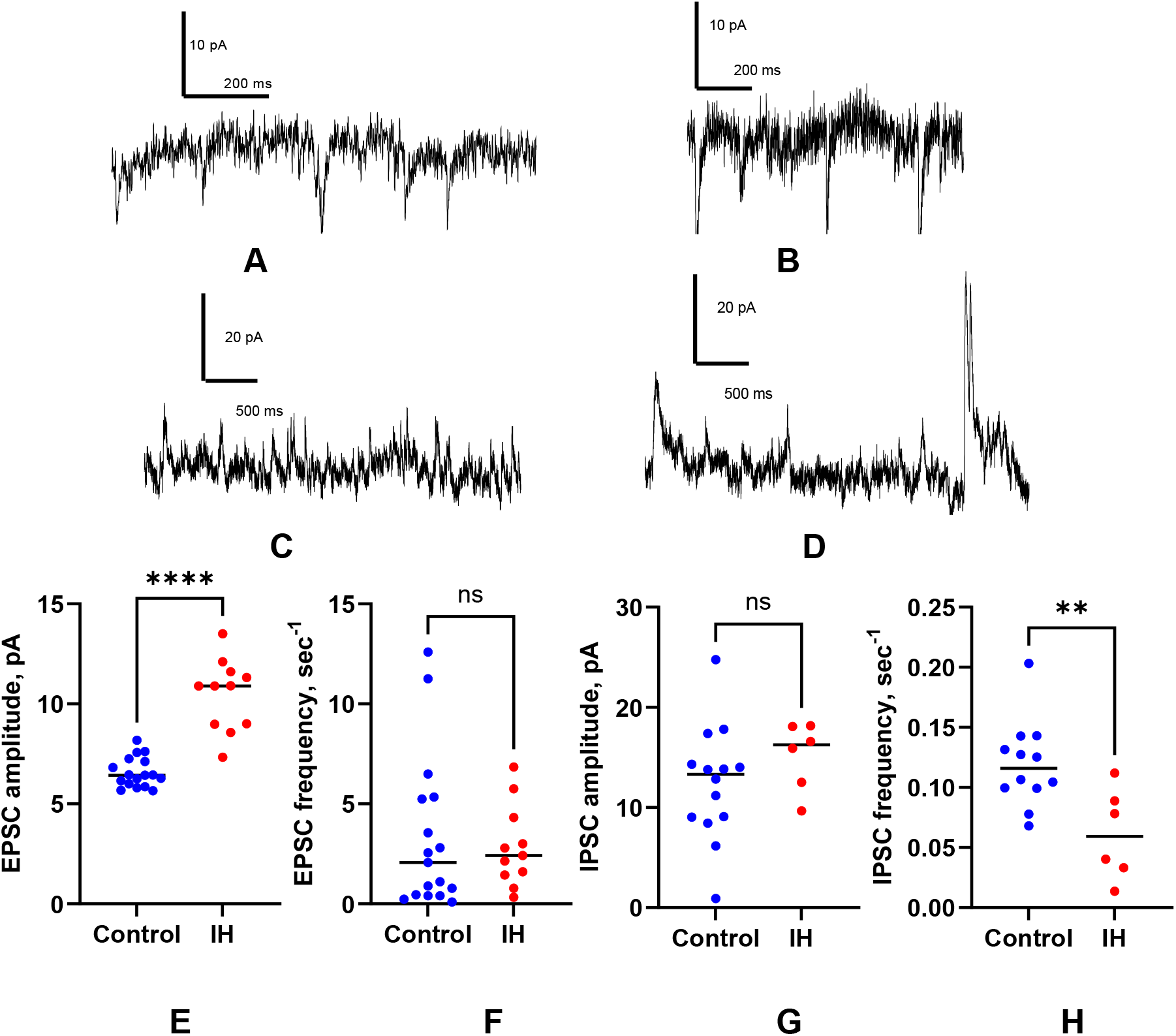
Patch clamp recording in pyramidal cells of primary motor cortex in adult mice after neonatal IH. A, B – examples of spontaneous glutamatergic postsynaptic currents recorded at a holding potential −70 mV in control (A) and IH (B). C, D - examples of spontaneous GABAergic postsynaptic currents recorded at a holding potential spontaneous 0 mV. EPSC’s amplitude (E), but not EPSC’s frequency (F) were increased in IH group. IPSC’s amplitude (G) was unchanged, but the frequency (H) was decreased in IH group **-p<0.01, ****-p<0.001, t-test

### Age-dependent effect of GABA inhibition on glutamatergic transmission in motor cortex

The sequence of field responses, consisting of the short latency volley burst (VB) followed by the first short latency fEPSP and often by the longer latency second fEPSP, was reproducible in all subgroups and observed in both control aCSF and after PTX application (Fig.3A). The VB was insensitive to neurotransmitter receptors blockade and was presumed to be presynaptic. The later components, first and second field potentials, were completely blocked by 50 µM CNQX and reflected monosynaptic (latency 4-6 ms) and polysynaptic activity (latency over 6-10 ms), both mediated by glutamate via AMPA receptor (Wallace, Jackson et al. 2014).

There were no other components in field potentials, such as NMDA kainate or GABAergic transmission components after CNQX and PTX block. VBs were completely removed by blocking axonal transmission with 1 µM of Tetrodotoxin (TTX). To ensure that the VBs were not affected by either age or PTX we compared VB amplitudes for all subgroups by single-factor ANOVA (F (31,234) =1.04, p=0.42). This comparison did not reveal a significant influence of either PTX. VB amplitudes increased linearly proportional to stimulation intensity in input-output curves. Since both criteria were met, we conclude that a comparison of the absolute values of FEPSPs can be used for the following analysis.

To examine the developmental change of GABA receptors inhibition on glutamatergic transmission we recorded fEPSPs in motor cortex slices before and after the addition of PTX to perfusate in pups between P7 and P13 when the developmental switch in GABA action in pyramidal neurons of cortical slices was expected (Peerboom and Wierenga 2021). PTX slightly decreased fEPSP between P7 and P8 (Figure 3B, Supplementary Figure 2), indicating excitatory action of GABA at this age. Starting at P9 in IH pups and at P10 in controls, blockage of GABA receptors by PTX increased the amplitude of fEPSP (main effect of age on two-factor ANOVA F (7, 66) = 13.02, p<0.0001), indicating a switch to inhibitory GABA action. The increase of fEPSP after PTX was more than twice as large in the IH group (main effect of IH, F (1, 66) = 9.700, p<0.0027) and significant between P9 and P13 (post hoc p<0.001), but not different in adults.

**Figure 3.**
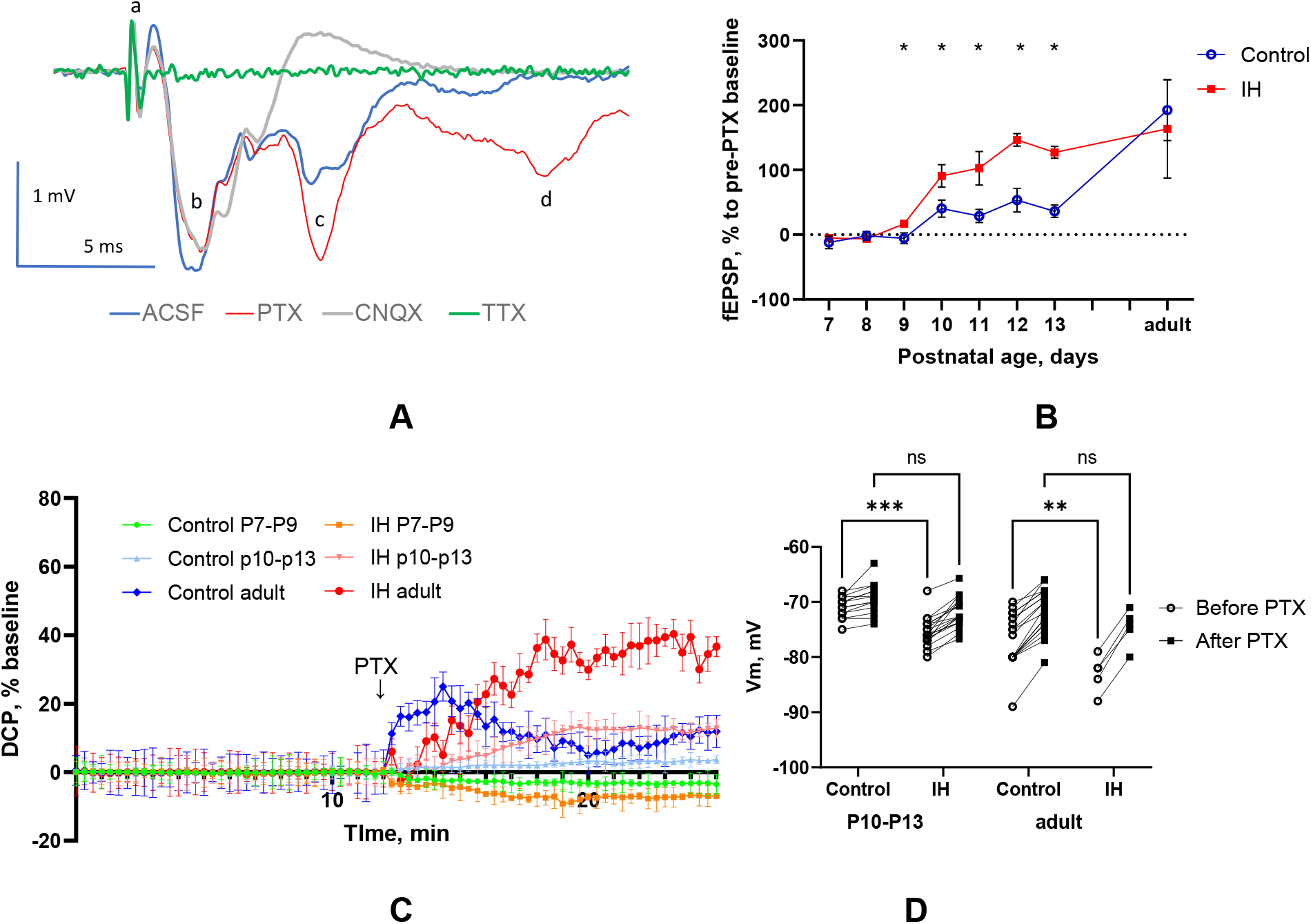
Effect of GABA inhibition on post-synaptic field potential in mouse motor cortex slices. A. Examples of fEPSP and waveforms origination at P8. a - truncated stimulation artefact, b – volley burst, c – first monosynaptic fEPSP, d-second polysynaptic fEPSP. Legend below shows inhibitors used to identify origin of components. B. Transient increase of monosynaptic fEPSP after PTX in IH group. C. Increase of direct current potential after PTX in IH group after P7. D. Transmembrane potential was increased after neonatal IH at P10-P13 and in adults, but could be returned to control values after PTX.

To investigate the effects of changes in tonic inhibition on glutamatergic transmission following IH, the resting direct current potential (DCP) was monitored at the beginning of each sweep before the stimulation artifact. Once a stable DCP was achieved for at least 15 minutes, PTX was applied. Since there was no spontaneous activity at the 200 µM concentration of PTX, the tonic change of DCP before stimulation after PTX application could be attributed to the blockage of extra-synaptic GABA receptors (GABA_A_R). The deflection of DCP from the baseline after 15 min of PTX application switched from negative at P7 to positive after P10 (Figure 3C) and was larger in the IH adult group than in control adults at 15 min after PTX (AVOVA F (5, 53) = 6.253, p=0.0001, post hoc p=0.0067).

Patch-clamp recordings of pyramidal cells (Figure 3D) confirmed an increase in tonic inhibition mediated by extrasynaptic GABAARs. This effect was observed in neonatal animals beyond P9 and persisted with a further increase in adult animals. The absolute values of Vm were larger (more negative) after IH both at P10-P13 and in adults (RM ANOVA F (3, 54) = 10.17, p=0.0001) than in age-matched controls. Vm values were reduced to those in the control groups at corresponding ages with the application of PTX (no difference between post-PTX groups on post-hoc comparisons), indicating that the increased hyperpolarization after IH was due to the activity of extrasynaptic GABA_A_R.

### Picrotoxin increased spontaneous MUA in IH but not in control mice in vivo

Following a low dose (0.1 mg/mL)) of PTX application on the cortex surface of the control group of mice, there was no significant change in MUA relative to the pre-PTX baseline, 99.25 ± 22.5% (Figure 4A, n = 6, t-test, ns). The mean increase after PTX injection was 213.74 ± 26.15% (Figure 4B, N = 5, t-test, p < 0.013) relative to pre-PTX baseline. Mean MUA change relative to baseline was higher in the IH groups than in the controls (Figure 5C, t-test, p < 0.009).

**Figure 4.**
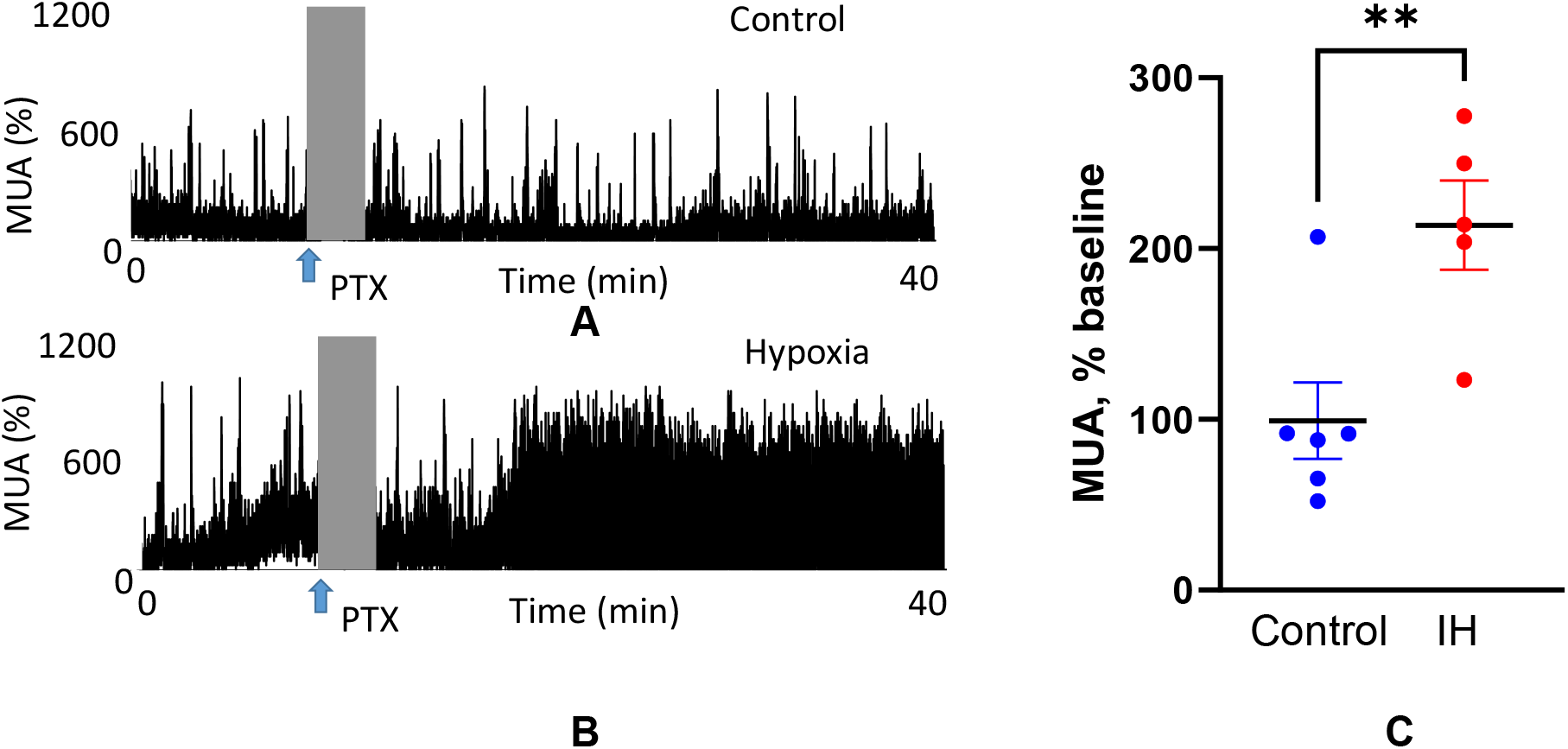
Examples and group data of extracellular multi-unit recording in motor cortex of awake neonatal mice at P12-P14. Control (A) and intermittent hypoxia (B) animals are shown. After 9 min of baseline recording picrotoxin 1 µL of PTX 0.1 mg/mL was superfused on exposed cortex above the recording electrode (arrow). C. MUA increase was significantly larger in IH group. **-p<0.01, t-test

**Figure 5.**
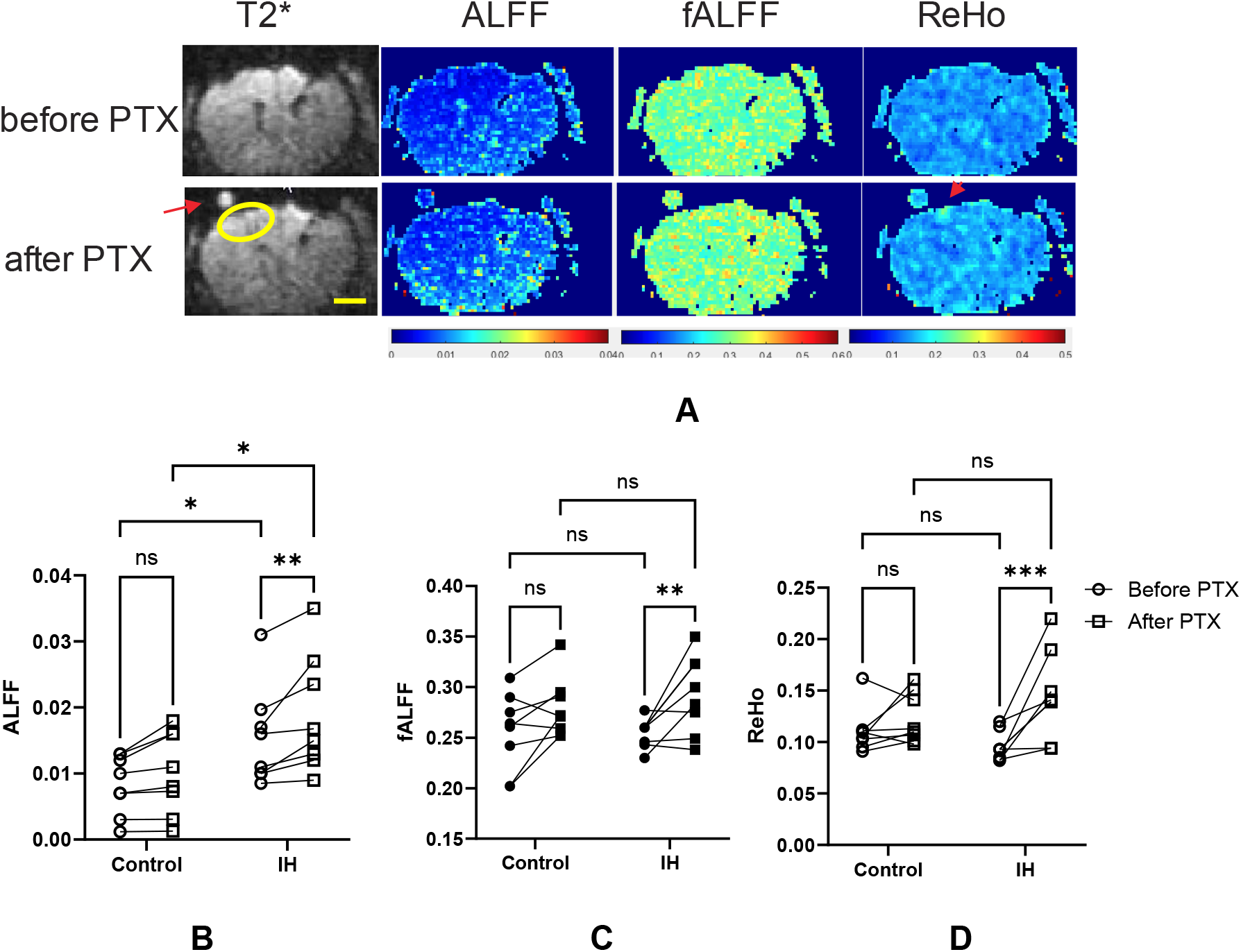
Changes in markers of cortical short range functional connectivity after pharmacological GABA inhibition on rsfMRI at P12. A. Representative T2* images and corresponding local functional connectivity indexes before and after cortical PTX injection in a mouse pup after IH. Arrow indicates a MRI visible marker placed on the site of PTX injection. Arrowhead points on the local increase of ReHo connectivity in the site of injection. Oval ROI was placed on the site of injection. Scale bar 1mm.B-D. Individual animal changes in markers of short range functional connectivity after ALFF, fALFF and ReHo in cortical region of interest around PTX injection.*-p<0.05, **-p<0.01, ***-p<0.0-1, RM ANOVA post-hoc.

### Increased ALFF and regional signal coherence in mouse pups after neonatal IH

Resting-state fMRI was performed on awake mouse pups at P12. Representative maps of local functional connectivity indexes before and after cortical PTX injection are shown in Figure 5A. A repeated measure two-way ANOVA was performed to evaluate the effects of IH and pharmacological GABA inhibition after PTX injection on indexes of local functional connectivity in the motor cortex. The results for ALFF indicated significant main effects for IH, F (1, 14) = 5.460, p = .034, and for PTX F (1, 14) = 17.36, p=0009. The interaction between IH and PTX was not significant. ALFF was higher in the IH group (p=0.04) before PTX. PTX injection increased ALFF in the IH group (p=0.0016), but not in the control group. PTX injection also increased fALFF (F (1, 14) = 14.45, p=0.0019, RM ANOVA, p=0.005 post hoc) and ReHo (F (1, 14) = 13.75, p=0.0023, RM ANOVA, p=0.0008 post hoc) in IH groups, but not in the control groups.

### Increased gene expression in markers of glutamatergic transmission but not in markers of interneuron function

To elucidate the molecular mechanisms underlying the observed changes in interneuron-mediated regulation of glutamatergic transmission and neuronal synchronization, we compared gene expression profiles in the motor cortex of control and IH pups. We focused on a predefined set of genes involved in ion channel function and synaptic transmission (Supplementary Figure 3). Our findings align with previous reports (Goussakov, Synowiec et al. 2021) by demonstrating a significant upregulation of several AMPA receptor subunits in the IH group, consistent with enhanced AMPA receptor-mediated transmission. Interestingly, we observed no significant changes in the expression of the GABA synthetic enzyme glutamate decarboxylase (GAD), including GAD1 and GAD2, which are considered reliable markers of GABA release (Dicken, Hughes et al. 2015)

Similarly, no significant differences were observed between control and IH groups at key developmental stages in the expression of Slc12a2 and Slc12a5, genes encoding NKCC1 and KCC2 proteins that regulate the shift of GABA from excitatory to inhibitory. This finding aligns with our field potential recordings using Picrotoxin (PTX), suggesting that IH does not affect the timing of the GABA switch. Interestingly, we observed a trend towards elevated expression of the delta subunit of GABA receptors in the IH group at postnatal days 14 and 30, Supplementary Figure 3), potentially linked to extra-synaptic tonic GABA inhibition, which aligns with our findings in the DCP and Vm at those ages (Figure 3C). However, this increase did not reach statistical significance using ANOVA.

### Long-term decrease in PV expression in the somatosensory cortex after neonatal IH

To examine changes in inhibitory interneurons, we quantified the density of major interneuron types expressing parvalbumin (PV+) and somatostatin (SST+) immunoreactivity in the somatosensory cortex of mice at P12, 5 days after exposure to neonatal intermittent hypoxia (IH) between P3 and P7, and at P60 when the brain is mature There was a significant decrease in the density of PV+ interneurons in the IH group compared to normoxic controls at P12 and at P60 (main effects for IH, F (1, 8) = 67.44, p <0.0001, post-hoc p=0.0045 for P12 and p=0.0008 for P40, Fig. 6A). The density of neurons expressing PV increased with age both in control and

**Figure 6.**
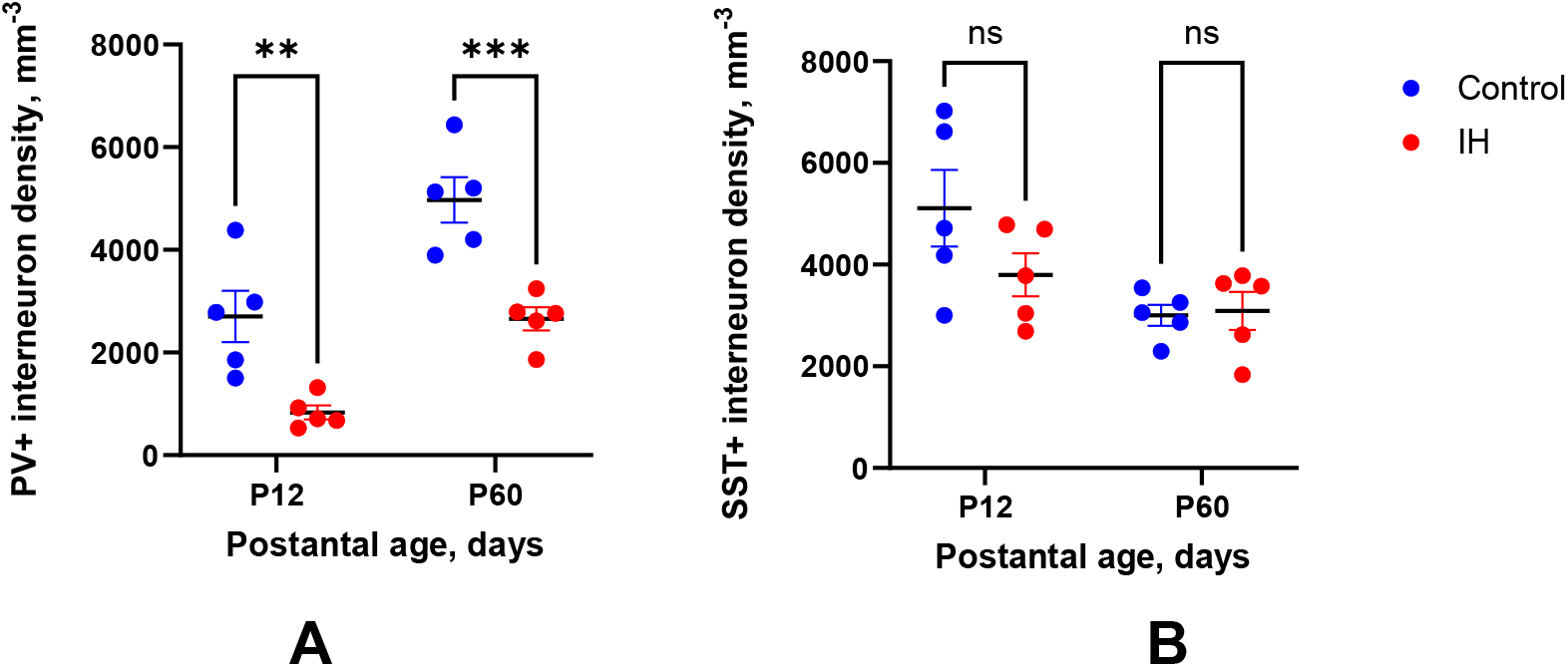
Stereological count of interneurons, labelled for parvalbumin (A) and somatostatin (B) in sensory-motor cortex at P13 after neonatal IH. **-p<0.01, ***-p<0.0-1, ANOVA post-hoc

IH groups (F (1, 8) = 21.17, p=0.0018). The interaction between IH and age was not significant. The difference between the groups in SST+ neuron density was not statistically significant for both ages (F (1, 8) = 1.601, ns, Fig. 6B). There was a decrease in the density of SST+ neurons in the older animals (F (1, 8) = 8.504, P=0.0194).

## DISCUSSION

We investigated the interaction between excitatory and inhibitory neurotransmission during a critical stage of cortical network development after perinatal HI brain injury. The major findings of the study were 1) increased glutamatergic transmission, evidenced by patch clamp miniature currents and field potential recordings with a selective GABA receptor antagonist picrotoxin, 2) the decreased inhibitory drive, based on lower IPSC frequency on pyramidal cells and MUA recordings with Picrotoxin injection, immediate and long-term decrease of PV+ interneuron density and 3) increased tonic inhibition by GABA_A_Rs and in the motor cortex 6 days after neonatal IH. The other significant result in this study was abnormally increased neuronal connectivity in the actively developing cortex at the second postnatal week, reflected by the elevated low-frequency fluctuations in resting-state fMRI in the IH group and further enhanced by Picrotoxin. Long-term behavioral consequences, locomotor hyperactivity, and impaired motor learning were observed in adult mice exposed to neonatal IH.

Perinatal hypoxic brain injury occurs at the critical period of cortex development and activity-dependent formation of functional neuronal networks (Blankenship and Feller 2010), governed by maturing excitatory and inhibitory signaling. We have previously reported abnormally increased excitatory glutamatergic AMPA and NMDA–mediated transmission in the hippocampus after perinatal IH (Goussakov, Synowiec et al. 2021) and now confirmed it in the motor cortex by the increased EPSC’s amplitude in principal cells. Maturation of the inhibitory GABAergic system occurs later in the development and the transition from excitatory to inhibitory action of GABA constitutes a major change in the pattern of neuronal activity (Ben-Ari, Gaiarsa et al. 2007). In our field potential recording with PTX during the first two postnatal weeks in mice, GABA transmission provided an excitatory effect before switching to inhibition at P9 in the IH group and P10 in the controls. There is evidence that the timing of the GABA switch is influenced by neuronal activity (Fiumelli, Cancedda et al. 2005)and may be affected by neonatal IH. Neonatal IH caused the GABA switch to occur one day earlier compared to controls. However, the relative gene expression of NKCC1 and KCC2 chloride transporters did not differ between the groups. Despite the minimal difference in timing, the acceleration of the GABA switch may still have significant consequences, potentially leading to impairments in motor learning and cognitive behavior, similar to what has been observed in mice with early life stress (Karst, Droogers et al. 2023) and other neurological disorders (Ben-Ari, Khalilov et al. 2012, Peerboom and Wierenga 2021).

After applying PTX at P10, the magnitude of the monosynaptic fEPSP increased in control mice. However, this increase was nearly double in the IH group. The larger fEPSP changes likely result from two contributing factors: the previously described increase in AMPA transmission and a larger, PTX-sensitive GABAergic inhibition. This inhibition was tonic since it was measured in the short-latency monosynaptic response, not involving either feedback or feed-forward inhibitory circuits. It does not involve phasic inhibition mediated by the synaptic release of GABA from interneurons, and it could be mediated by extra-synaptic GABA receptors (Marchionni, Omrani et al. 2007). Further confirmation of increased tonic GABA_A_R inhibition after neonatal IH was obtained by recording DCP and Vm before and after PTX application.

Consistent with the age-related changes in fEPSP, the deflection of DCP from negative to positive occurred around P9-P10 and was larger in the IH group, in both P10-P13 pups and adults. Transmembrane potential due to the activity of extra-synaptic GABA_A_R was increased within a few days after IH and in adults. The increase in tonic inhibition mediated by extrasynaptic GABAARs might be a compensatory response to the upregulation of AMPA receptors and heightened glutamatergic activity after neonatal IH.

GABAergic interneurons are the major population of cortical inhibitory neurons, representing approximately 20% of all cortical neurons (Sahara, Yanagawa et al. 2012). The effect of perinatal hypoxia on interneuron development remains controversial. An increase in the density of the perineuronal nets and PV+ interneurons was observed in adult rats treated with moderate hypoxia on the first day of life (Trnski, Nikolić et al. 2022). On the other hand, the numbers of cortical interneurons expressing immunohistochemically detectable levels of PV and SST, and vasoactive intestinal peptide were decreased by P15 in hypoxic-reared mice (Komitova, Xenos et al. 2013). Since the number of GAD+ neurons did not change, the authors suggested a delay in maturation, but not death of PV + interneurons after hypoxia. However, a loss of GAD+ and PV+ interneurons and disruption of perineuronal nets was found in the cerebral cortex following hypoxia-ischemia in near-term fetal sheep (Fowke, Galinsky et al. 2018), at P18 in mice hippocampus after neonatal hypoxia-ischemia (Chavez-Valdez, Emerson et al. 2018), in inflammation-induced perinatal injury in mice and humans (Stolp, Fleiss et al. 2019). The timing and severity of hypoxic insult may affect the numbers and maturation of interneurons in the developing cortex. Our study revealed a decrease in the inhibitory drive due to an immediate and long-term decrease of PV+ (but not SST+) interneuron density and functional interneuron deficiency by decreased IPSC frequency. Considering that expression of PV and maturation of PV+ neuron firing properties depends on the excitatory synaptic drive (Miyamae, Chen et al. 2017, Warm, Schroer et al. 2021), it is not clear why the decreased inhibitory drive coincides with the elevated glutamatergic transmission in the cortex after perinatal hypoxic brain injury and requires further investigation.

In the current study, we demonstrate for the first time the functional deficiency of cortical interneurons after perinatal brain injury by recording on non-anesthetized neonatal animals with the application of a GABA blocker. Applying a low dose of PTX did not affect multi-unit activity (MUA) in control animals. However, it significantly increased MUA in IH animals. This suggests a reduced capacity of the GABAergic inhibitory system to control excessive excitation and prevent hyper-synchronization of neuronal activity in the IH group. The observed functional deficiency in the inhibitory system and the increased glutamatergic transmission likely disrupt neuronal network formation, contributing to hyperactivity and cognitive deficits (Goussakov, Synowiec et al. 2021). Locomotor hyperactivity is a well-established finding after perinatal hypoxia (Mikati, Zeinieh et al. 2005, Trnski, Nikolić et al. 2022), and this study confirmed it in adult female mice. However, deficits in motor learning were more subtle and only detectable using a complex wheel task. By ensuring equal active time in the running wheel between groups, we were able to isolate motivational and hyperactivity differences from the specific motor learning deficiency.

Our finding of coexisting increased tonic inhibition with decreased synaptic inhibition highlights the complex regulation of the excitation/inhibition balance in the developing brain after injury. This interplay involves phasic, synaptic, and GABAergic modulation of glutamatergic transmission. Interestingly, despite the decrease in synaptic inhibition, the overall effect of neonatal hypoxia on neuronal circuitry appears to be excitatory, as shown in the current and previous data (Jensen, Wang et al. 1998, Goussakov, Synowiec et al. 2021). A critical implication is that a substantial rise in excitatory transmission alone can lead to completely dysfunctional networks prone to seizures. Therefore, compensatory mechanisms, such as the observed robust elevation of tonic inhibition directly targeting monosynaptic glutamatergic inputs, become essential for maintaining network functionality even during heightened excitatory drive.

The ability to non-invasively assess the functional development of somatosensory cortical networks after perinatal hypoxia is of significant clinical importance. Here, we investigate the potential of resting-state functional magnetic resonance imaging (rs-fMRI) to assess functional connectivity within this brain region. We hypothesized that disturbances in the inhibitory system maturation could be reflected in the pattern of local blood flow fluctuations, likely due to both synchronized neuronal activity and the direct role of interneurons in regulating vasomotion (Attwell, Buchan et al. 2010). Local functional connectivity, measured by rsfMRI, is likely depends on the activity and architecture of a small-scale network of primary cells glutamatergic neurons, GABAergic interneurons of various types, astrocytes, and micro-vessels, where the BOLD signal originates. However, the mechanisms of those interactions and the contribution of specific cell types remain poorly understood. Short-range functional rsfMRI connectivity has been described to increase in fetuses and neonates (Cao, He et al. 2017, Huang, Wang et al. 2020) and then decrease later into adulthood (Ouyang, Kang et al. 2017), depending on the anatomical region (He and Parikh 2016). The inverted U-shaped trend is often attributed to a combination of synaptogenesis followed by synaptic pruning, although the exact mechanisms remain unclear. Atypical development of local rsfMRI connectivity is associated with major cognitive and behavioral disorders, including ADHD (Sato, Hoexter et al. 2012, Sripada, Kessler et al. 2014), schizophrenia and autism (Fair, Cohen et al. 2009).

The absolute increase of ALFF in the IH group of this study suggests an increased level of synchronization in neuronal activity with a corresponding effect on blood flow in neighboring cortical voxels. The contribution of interneurons to the synchronization process was confirmed by an increase in all reported indexes of local functional connectivity after PTX injection. A larger increase of ALFF and ReHo after PTX relative to controls suggests that PTX revealed an interneuron dysfunction in the IH group. This dysfunction was not obvious without PTX injection, perhaps due to the effective compensatory abilities of local neuronal networks. A higher regional ALFF was found in infants born prematurely (Feng, Wang et al. 2024) and was associated with higher levels of whole-brain connectivity in children with autism (Supekar, Uddin et al. 2013) and in certain areas in children with attention deficit hyperactivity disorder (Yu-Feng, Yong et al. 2007). An absolute increase in fALFF and a relatively large increase of fALFF after PTX in the IH group indicated that the spectral power of the BOLD signal is shifted to an area of lower frequencies, and this shift has the contribution of interneurons. In a human study in 1-year-old infants, a right-ward shift of BOLD signal frequency was found during the first year of life, and the power of peak frequency in the sensorimotor and lateral visual networks showed positive correlations with motor and visual performance (Alcauter, Lin et al. 2015). Understanding the relationship between rsfMRI measures and functional cortical development will be critical for establishing this technique as a valuable tool for evaluating therapeutic interventions in neonates with hypoxic brain injury.

Overall this study demonstrates that perinatal brain injury disrupts the balance between excitatory and inhibitory neurotransmission in developing cortical networks. This disruption, potentially caused by functional deficiencies in GABAergic interneurons alongside increased glutamatergic transmission, may contribute to altered brain connectivity and the observed behavioral deficits, including hyperactivity and cognitive difficulties. This study underscores the potential of fMRI as a valuable clinical tool to elucidate mechanisms of functional maturation and abnormal connectivity in the developing cortex following perinatal hypoxic brain injury.

## Supporting information

Manuscript

## FUNDING

This study was funded by NIH grants 1R01NS119251-01A1, R01 NS107383, R01 GM112715, S10OD032223

## Code and data availability statement

The data used in this study are included in the manuscript and supplementary data.

## Authorship contribution statement

Ivan Goussakov, Alexander Drobyshevsky - conceptualization, data acquisition, analysis, writing Sylvia Synowiec, Rafael Fabres, Daniil Aksenov, Gabriela Almeida, Silvia Takada – data acquisition and processing

## Notes

### Competing Interest Statement

The authors have declared no competing interest.

