## Supplementary material for "Abnormal local cortical functional connectivity due to interneuron dysmaturation after neonatal intermittent hypoxia": Manuscript

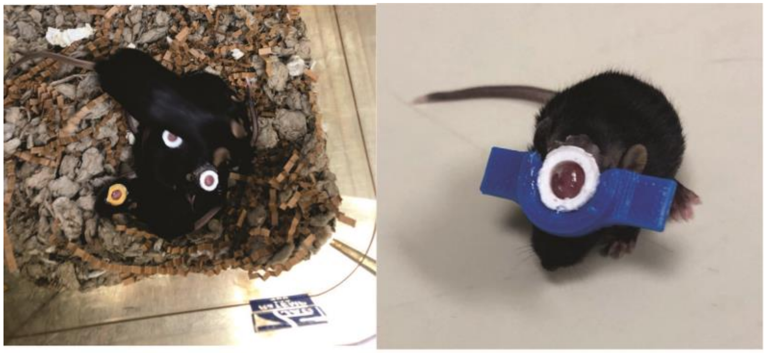

A B

Supplementary Figure 1. Neonatal mice instrumentation for in vivo MRI and electrophysiological recording. A. At P11 mouse pups were implanted with a custom 3D printed circular head post 10 mm diameter and opening 6 mm diameter with a small height profile to minimize disturbances to the dam while nursing in nest. B. During MR imaging and electrophysiological recording the head post was coupled with an adapter (blue) allowing head fixation to the animal cradle.

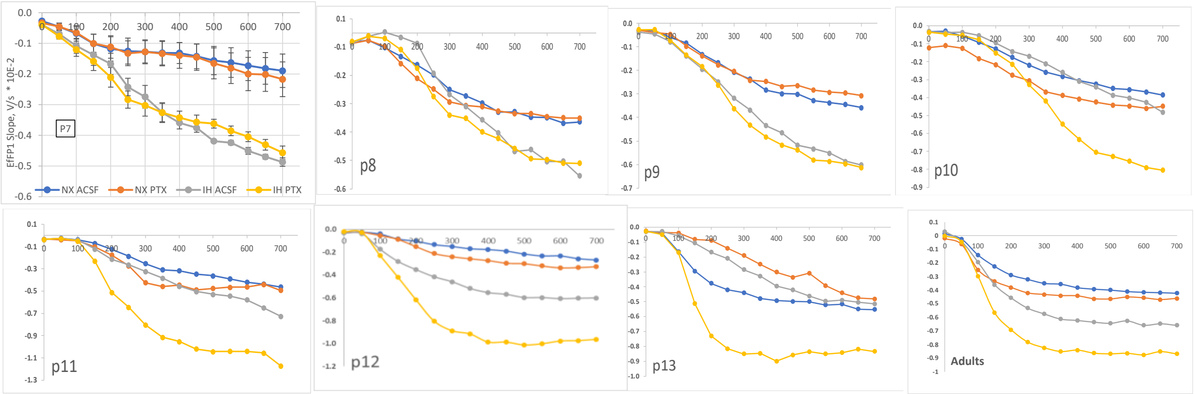


Supplementary Figure 2. Input –output curves recorded in neonatal mice between P7 – P13 and in adult at 50 days old in control normoxic group (NX) and after neonatal intermittent hypoxia (IH). The I-O curve were recorded in aCSF before and after addition of 200 µM of Picrotoxin (PTX). The slope of the first fEPSP is show on y-axis.

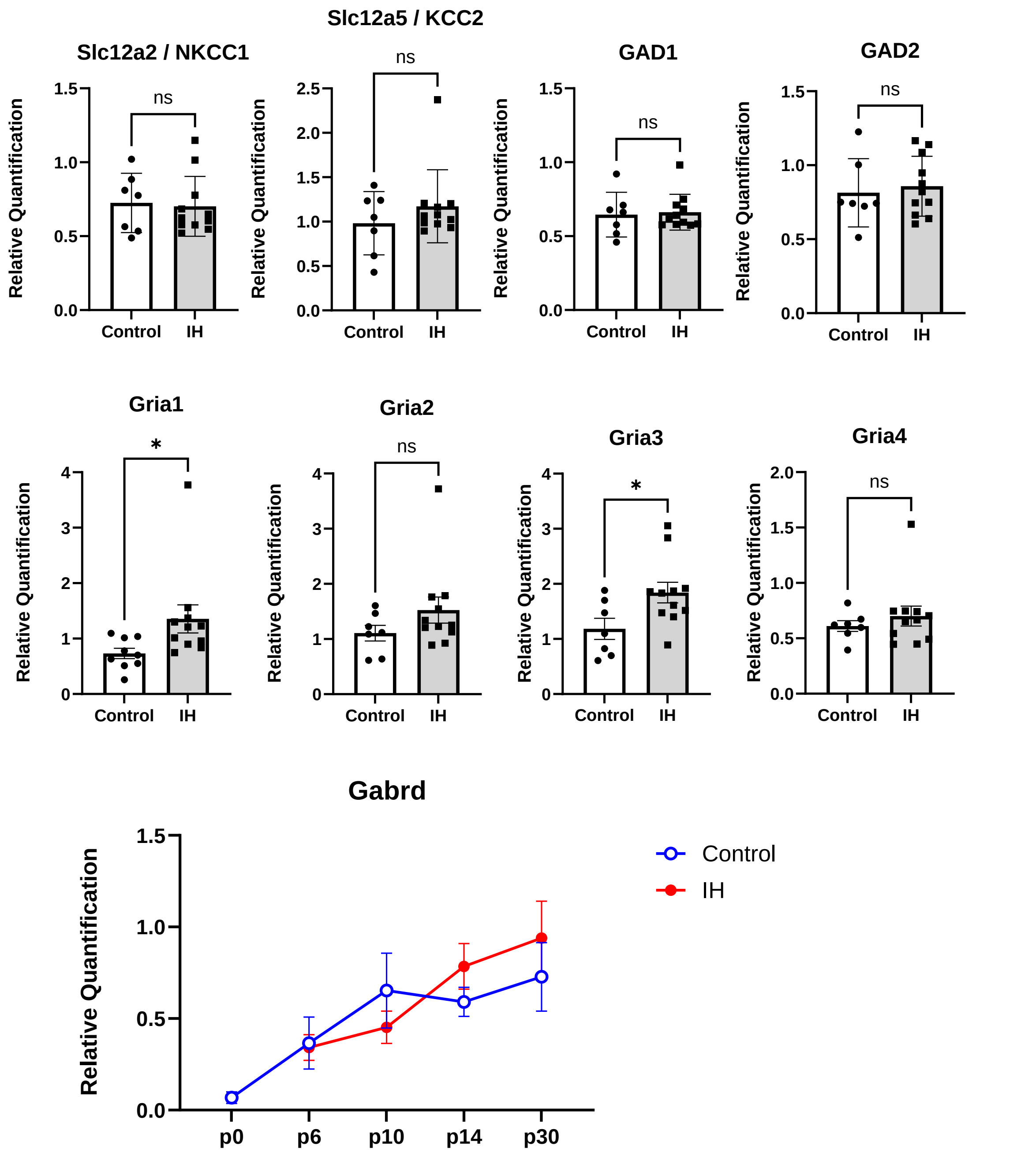


Supplementary Figure 3. Relative gene expression of ion channels and interneuron function after neonatal IH. Mean ±SEM. * - p<0.05, two-tailed t-test

Supplementary Table 1. Primers used in the study.

| Primer | Foward | Reverse | Primers designed |
|---|---|---|---|
| GAPDH | AAT GGT GAA GGT CGG TGT G | GTG GAG TCA TAC TGG AAC ATG TAG | IDT (NM_008084) |

| | | | |
|---|---|---|---|
| GRIA1 | ATC GAG TTC TGC TAC AAA TCC C | TCC GTA TGG CTT CAT TGA TGG | IDT (NM_008165) |
| GRIA2 | CAC TTC GGA GTT CAG ACT GAC | AAT CGC ATA GAC GCC TCT TG | IDT(NM_013540) |
| GRIA3 | GTG TGA TAC GAT GAA AGT TGG TG | AGA TGC CTT GTT CAC TGA GTT | IDT(NM_016886) |
| GRIA4 | GGT ACG ATA AAG GTG AAT GTG GA | CCG CCA ACC AGA ATG TAG AAG | IDT(NM_019691) |
| GAD1 | GGA CAT CTT CAA GTT CTG GCT | CTT GGC GTA GAG GTA ATC AGC | IDT(NM_008077) |
| GAD2 | GCA CTA TGA CCT GTC CTA TGA | GTG CCT CAA ACC CAG TAG TC | IDT(NM_008078) |
| SLC12a2 | ATC CTC AGT CAG CCA TAC | TCT GTG GGT TCG TGT GTT | This study |
| SLC12a5 | ACA ATG TCA CAG AGA TCC | GAG TTC TTA CCT GAC CAA | This study |
